# Regulation of B cell function and expression of CD11c, T-bet, and FcRL5 in response to different activation signals

**DOI:** 10.1101/2023.03.08.531830

**Authors:** Linn Kleberg, Alan-Dine Courey-Ghaouzi, Maximilian Julius Lautenbach, Anna Färnert, Christopher Sundling

## Abstract

CD11c, FcRL5, or T-bet are commonly expressed by B cells expanding during inflammation, where they can make up >30% of mature B cells. However, the association between the proteins and differentiation and function in the host response remain largely unclear. We have assessed the co-expression of CD11c, T-bet and FcRL5 in an in vitro B cell culture system to determine how stimulation via the B cell receptor (BCR), toll-like receptor 9 (TLR9), and different cytokines influence CD11c, T-bet, and FcRL5 expression. We observed different expression dynamics for all markers, but a largely overlapping regulation of CD11c and FcRL5 in response to BCR and TLR9 activation, while T-bet was strongly dependent on IFN-γ signalling. Investigating plasma cell differentiation and antigen-presenting cell (APC) functions, there was no association between marker expression and antibody secretion or T cell help. Rather the functions were associated with TLR9-signalling and B cell-derived IL-6 production, respectively. These results suggest that the expression of CD11c, FcRL5, and T-bet and plasma cell differentiation and improved APC functions occur in parallel and are regulated by similar activation signals, but that they are not interdependent.

## Introduction

B cells are important contributors to long-lived immunity through their differentiation to antibody-secreting plasma cells and long-lived memory B cells. This process is initiated in secondary lymphoid organs where antigen-activated mature naïve B cells either directly differentiate to plasma cells (T-cell-independent) in response to high-valent antigen or strong pattern-recognition-receptor (PRR)-signal, or migrate to T cell-B cell interfollicular to interact with activated cognate CD4^+^ T cells (T-cell dependent response) (1). In the T-cell-dependent response, B cells present peptide antigens to the cognate T cells via the major histocompatibility complex (MHC) class 2. Both the T cells and B cells then deliver additional activation and regulatory signals to each other via co-stimulatory molecules (such as CD40-CD40L, ICOS-ICOSL, CD80/86-CD28) and cytokines (2). At this stage, B cells can undergo cytokine-dependent class-switch recombination (3), enter the B cell follicle and establish germinal centres to undergo isotype- and affinity-dependent selection (4–6), eventually producing high-affinity memory B cells and plasma cells (7), or they can directly differentiate to relatively low-affinity extrafollicular memory B cells and plasma cells (8).

For certain infectious diseases with a strong or prolonged inflammatory component, such as malaria and HIV, the process of generating long-lived B cell-mediated immunity has been suggested to be inefficient (9). This has in part been associated with the expansion of a B cell subset lacking the expression of CD21 and CD27 while upregulating CD19 and CD20, the integrin CD11c, the transcription factor T-bet and the IgG receptor FcRL5, in addition to numerous other receptors associated with cell regulation and migration (10–12). These cells have been suggested to be exhausted or dysfunctional, in part due to reduced responsiveness to BCR-ligation and upregulation of several inhibitory receptors (12,13). However, a recent study by Ambegaonkar et al. suggests it could have more to do with how the antigen is presented (14), and it remains unclear what function and purpose these cells play in the immune response (15). Some studies have suggested that the cells could be primed to migrate to sites of infection or inflammation to provide localized differentiation to plasma cells or to provide antigen-presenting functions to local CD4^+^ T cells (15–17). In support of such functions, CD21^−^CD27^−^ B cells in malaria display an upregulation of the plasma cell-associated genes *PRDM1* and *CD38* and differentiate into antibody-secreting cells following interaction with T-follicular helper (T_FH_) cells (18). Additionally, mouse and human studies have indicated more efficient antigen uptake and upregulation of MHC2 and co-stimulatory molecules, potentially translating to an enhanced capacity to drive CD4^+^ T cell responses (14,18,19).

It has become clear that CD11c^+^ or T-bet^+^ B cells are not necessarily only associated with severe inflammatory disease, as they are found at low levels during steady-state that then increase with normal ageing of the host (20–22). These levels are, however, greatly expanded in many different infections and following vaccination (23). Additionally, cells with a similar phenotype are also expanded in several different autoimmune diseases that have an inflammatory component (24). Despite sharing many phenotypic and transcriptomic characteristics between different types of diseases (25), there are also differences that appear disease-specific, such as the expression of FcRL4 in HIV infection and FcRL5 in malaria (12,26). An additional source of variation could come from how the cells are defined in different studies, where varying combinations of IgD, CD21, CD27, CD11c, CD85j, T-bet, CXCR3, and FcRL5/FcRL4 have been used. This has also led to the cells having been given many different names, such as atypical B cells, age-associated B cells, exhausted B cells, tissue-like B cells, double negative (DN) and DN2 B cells (10,12,26–32). We have previously shown that the co-expression of several of these markers is changing over time after an acute malaria episode (27). It is therefore possible that they could represent different stages of differentiation or alternatively that the cells have received different stimuli prior to their differentiation.

In this study, we have investigated the regulation of CD11c, T-bet, and FCRL5, three of the most used markers of these alternative lineage B cells. We have used an in vitro culture system to determine how stimulation via the BCR, TLR9 or the response to different cytokines influences their expression. Additionally, we have assessed their functional properties, including differentiation to antibody-secreting cells and B cell-dependent CD4^+^ T cell help.

## Results

### Differential kinetics of CD11c, T-bet, and FcRL5 after stimulation

To better understand the regulation of CD11c, T-bet, and FcRL5 in B cells, we adapted an in vitro culture system described by Ambegaonkar et al. (33) to generate B cells with an atypical phenotype. We sorted total, naïve, and memory B cells and labelled them with cell-trace violet (CTV), enabling tracking of cell division. We then incubated the cells over 4-6 days in the presence or absence of anti-Ig (BCR ligation), CpG-C (TLR9-ligation), and the proinflammatory cytokine IFN-γ (**Figure 1A**). We measured cell expression of CD11c, T-bet, FcRL5, and CTV by flow cytometry. Expression of CD11c, T-bet, and FcRL5 were all significantly upregulated in total B cells after 2 days of stimulation but showed slightly different kinetics, where T-bet levels peaked at 2 days, followed by FcRL5 levels at 4 days and CD11c levels at 6 days (**Figure 1B**). Upregulation of the markers was not dependent on cell division, which was only observed after 4 days of stimulation (**Figure 1C**). Naïve and memory B cells showed similar kinetics as total B cells, although CD11c was significantly more upregulated on memory cells compared with naïve cells, while T-bet and FcRL5 expression overlapped (**Figure 1D**). Proliferation was slightly more pronounced for naïve cells, although there was a large variation between donors (**Figure 1E**). In summary, B cells stimulated with anti-Ig, CpG-C, and IFN-γ upregulated CD11c, T-bet, and FcRL5, although with different expression kinetics.

**Figure 1.**
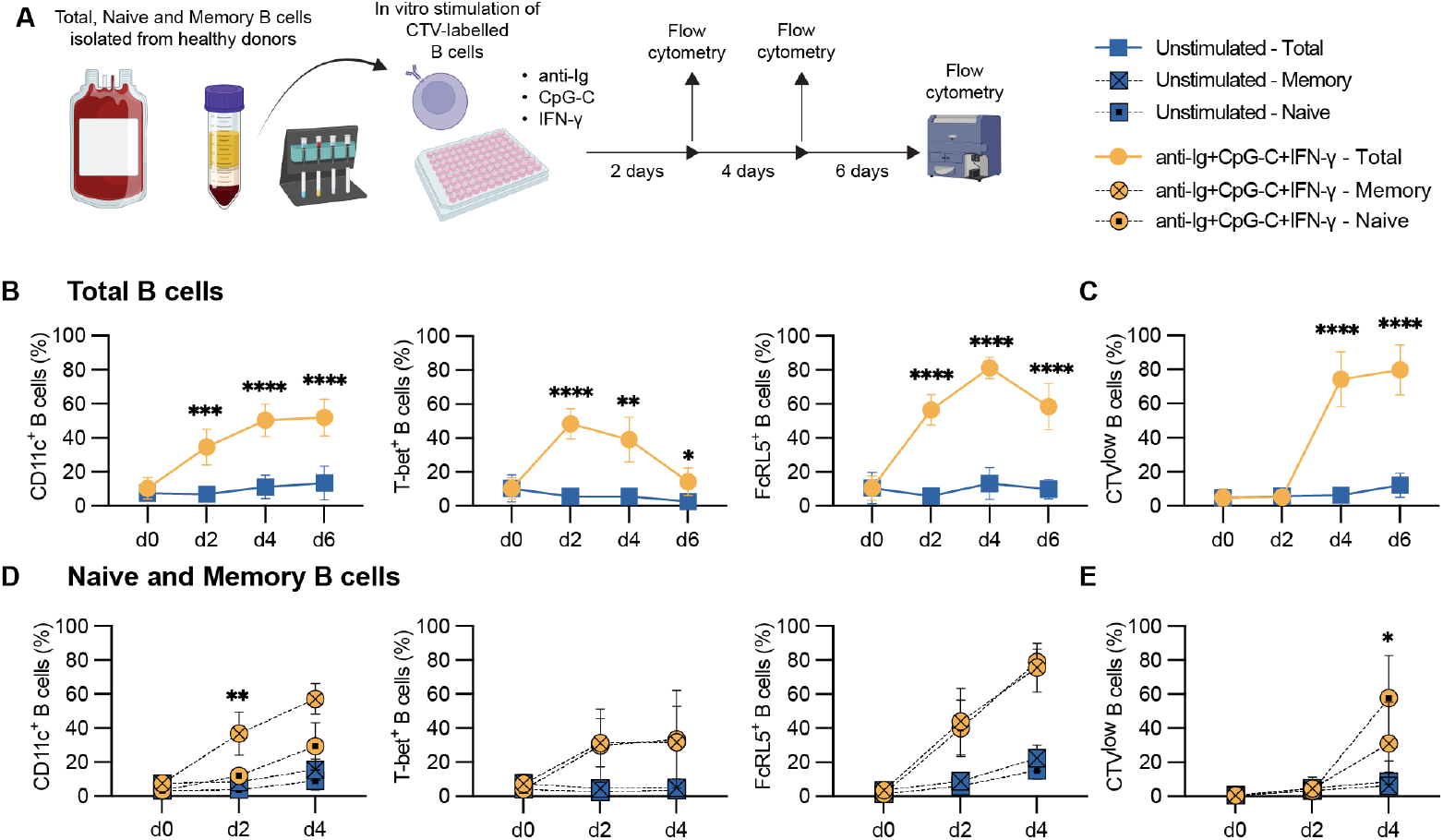
In vitro kinetics of B cell-expression of CD11c, T-bet, and FcRL5. (**A**) Schematic representation of experimental setup with B cells sorted from healthy blood donors using RosetteSep, followed by additional sorting into naïve and memory B cells using Magnetic beads. The cells were then stimulated with anti-Ig, CpG-C, and IFN-γ for 0, 2 and 4 days for naïve and memory B cells and for 0, 2, 4, and 6 days for total B cells. (**B**) Total B cell expression of CD11c, T-bet, and FcRL5 and (**C**) cell proliferation as indicated by the frequency of cell-trace violet low (CTV^low^) cells (n=6-9 donors). (**D**) naïve and memory B cell expression of CD11c, T-bet, and FcRL5 and (**E**) frequency of CTV^low^ cells (n=7 donors). Data is shown as the mean with a 95% confidence interval and is pooled from 3-4 separate experiments. Two-way ANOVA with Geisser-Greenhouse correction followed by Sidak’s posttest or in the case of missing data, by mixed-effects analysis were used for comparisons between stimulated and unstimulated in B-C and between naïve and memory B cells in D-E. No star indicates p > 0.05, *p<0.05, **p<0.01, ***p<0.001, ****p<0.0001.

### Role of BCR- and TLR9-ligation on B cell levels of CD19, CD11c, T-bet, and FcRL5

To evaluate the specific influence of BCR- and TLR9-ligation on cell marker expression, stimulation conditions without anti-Ig or CpG-C, in combination with IFN-γ, were compared to the condition with all three signals in sorted B cells from healthy blood donors (**Figure 2 and Supplementary table 1**). CpG-C was shown to be critical in inducing CD19^hi^ cells. On day 2, the removal of either anti-Ig or CpG-C led to a significant decrease in CD11c levels whereas, on day 6, only the condition without anti-Ig had significantly fewer CD11c^+^ cells. For T-bet levels, the addition of anti-Ig or CpG-C led to a significant increase of positive cells on day 2. On day 6, however, T-bet was down-regulated to baseline levels in all conditions. Both anti-Ig and CpG-C contributed to the up-regulation of FcRL5 on day 2, but no significant difference between stimulated and unstimulated cells was observed on day 6. In summary, TLR9-ligation was required for the upregulation of all four markers assessed here, while BCR-ligation led to up-regulation of CD11c, T-bet, and FcRL5, but not CD19.

**Figure 2.**
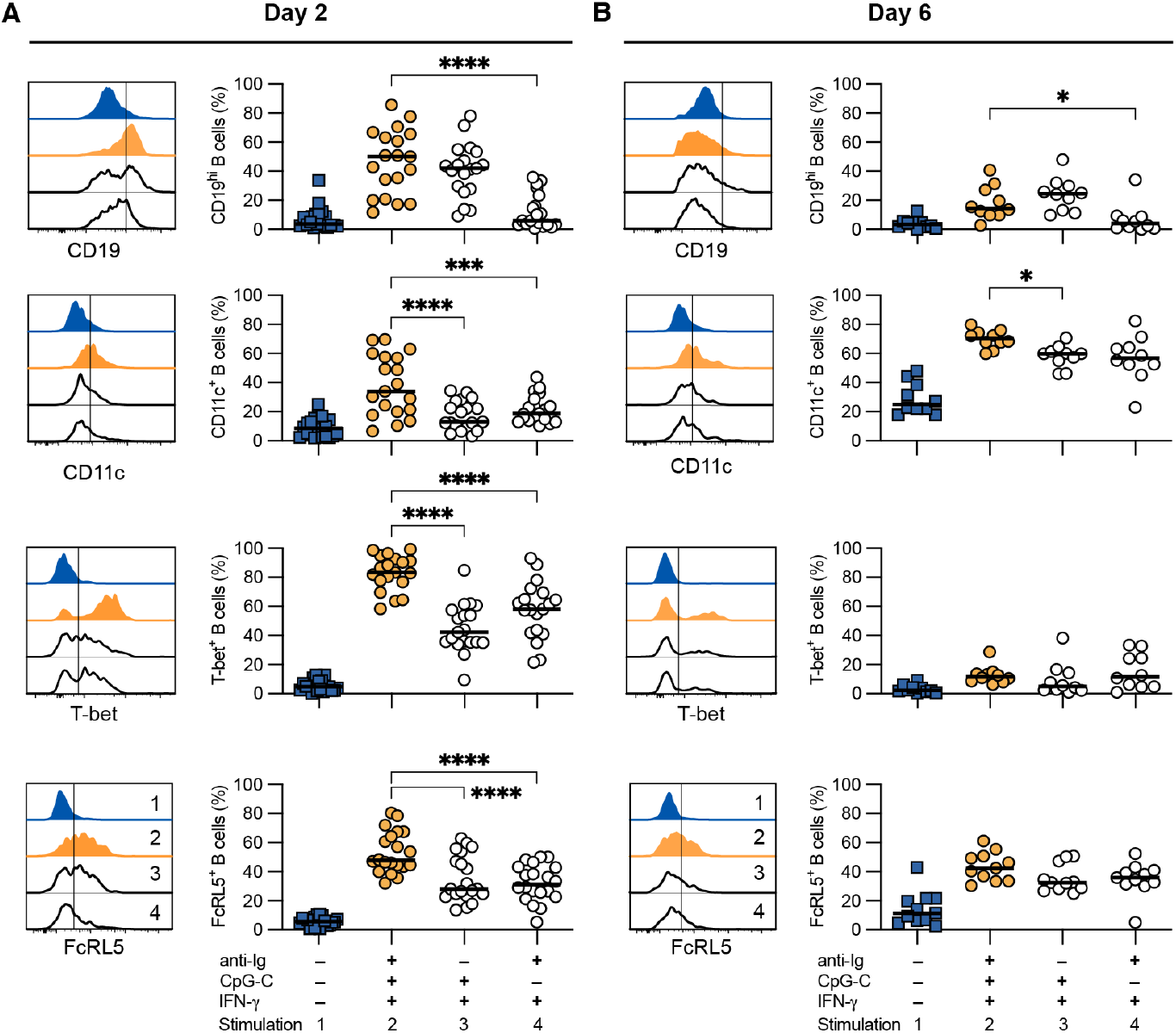
Role of BCR- and TLR9-ligation on CD19, CD11c, T-bet, and FcRL5 expression. Sorted total B cells from buffy coats were stimulated with combinations of anti-Ig, CpG-C, and IFN-γ for (**A**) 2 days (n=19-20) and (**B**) 6 days (n=10-11) after which expression of CD19, CD11c, T-bet, and FcRL5 was measured by flow cytometry. Representative histograms indicate marker expression. The line indicates the cut-off for positive expression. Numbers indicate the stimulations (see Supplementary table 1). Data is pooled from 9-17 separate experiments. P-values were calculated using matched pair one-way ANOVA with Geisser-Greenhouse correction followed by Sidak’s post hoc test. Unstimulated cells (blue boxes) were included for visual reference. Stimulated groups (open circles, stim 3 and 4) were compared with anti-Ig, CpG-C, IFN-γ (orange circles) stimulated cells. *p<0.05, ***p<0.001, ****p<0.0001. The absence of * indicates p>0.05.

### Differential impact of cytokines on B cell CD19, CD11c, T-bet, and FcRL5 levels

To further dissect the impact of the cytokines on the levels of CD19, CD11c, T-bet, and FcRL5, B cells sorted from healthy blood donors were cultured with anti-Ig, CpG-C, and either IFN-γ, IL-21, or IL-10 (**Figure 3 and Supplementary table 1**). None of the tested cytokines led to the upregulation of CD19 or FcRL5. The addition of IFN-γ resulted in a significant increase in CD11c levels, but only on day 2, while stimulation with IL-21 led to a significant increase on both days 2 and 6. IL-10 on the other hand had no significant effect on CD11c, although the mean CD11c^+^ frequency was somewhat higher than anti-Ig and CpG-C alone on day 2. In contrast, IL-10 stimulation led to a reduction of T-bet^+^ cells compared with anti-Ig and CpG-C stimulated cells alone. Only IFN-γ out of the 3 cytokines resulted in a significant increase in T-bet^+^ B cells on day 2, and a slightly higher percentage of cells remaining on day 6. This effect was not adversely affected by IL-10 (**Supplementary Figure 1**). In summary, T-bet expression was highly dependent on IFN-γ, while CD11c was upregulated in response to both IFN-γ and IL-21. CD19 and FcRL5 on the other hand were not influenced by the tested cytokines.

**Figure 3.**
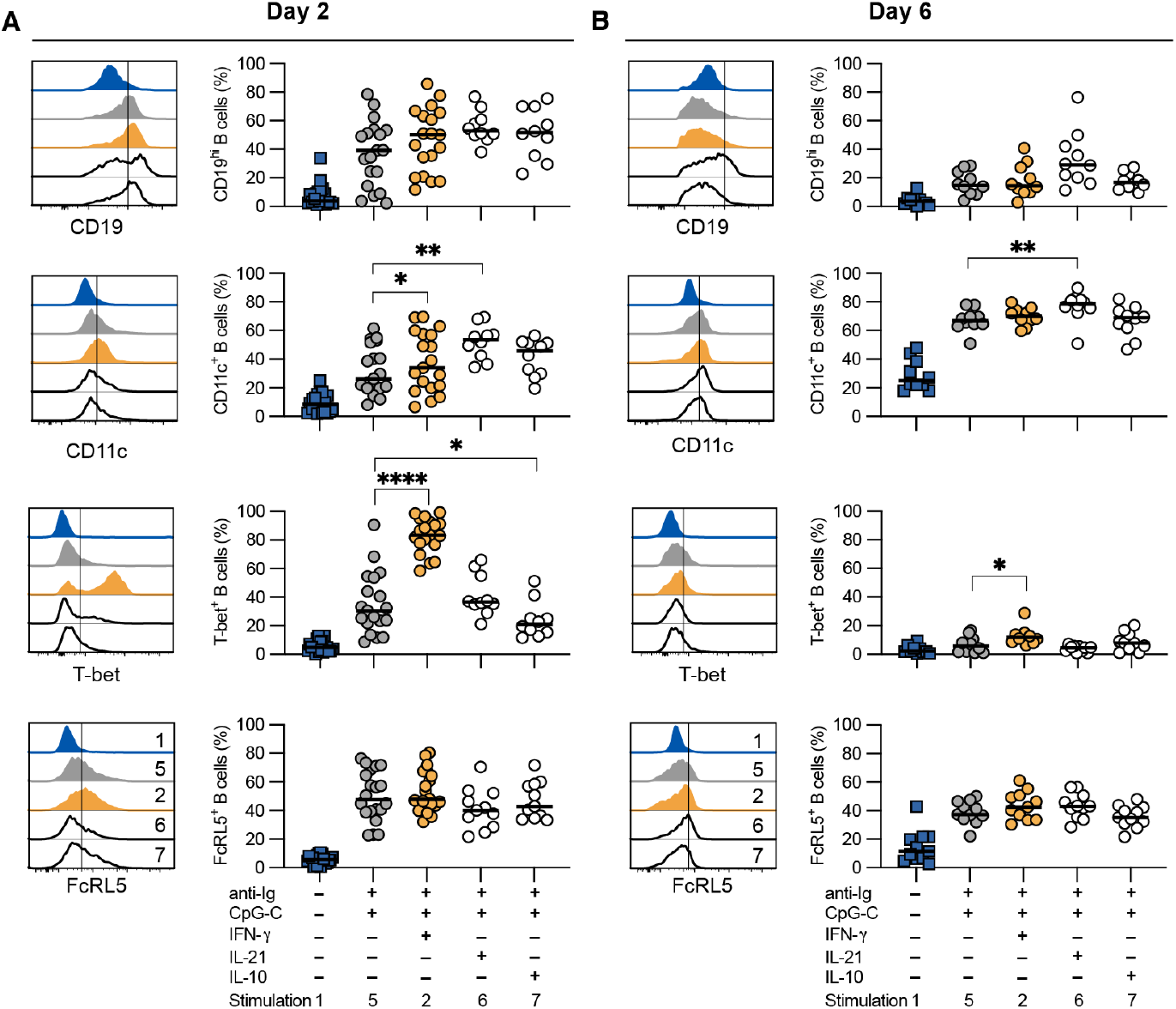
Role of IFN-γ, IL-21, and IL-10 on CD19, CD11c, T-bet, and FcRL5 expression. Sorted total B cells from buffy coats were stimulated with combinations of anti-Ig, CpG-C, IFN-γ, IL-21, and IL-10 for (**A**) 2 days (n=10-20) and (**B**) 6 days (n=10-11) after which expression of CD19, CD11c, T-bet, and FcRL5 was measured by flow cytometry. Representative histograms indicate marker expression. The line indicates the cut-off for positive expression. Numbers indicate the stimulations. Data is pooled from 9-17 separate experiments. P-values were calculated using matched pair one-way ANOVA with Geisser-Greenhouse correction followed by Sidak’s post hoc test or in the case of missing data, by mixed-effects analysis. Unstimulated cells (blue boxes) were included for visual reference. All stimulated groups had received anti-Ig and CpG-C. Stimulations 2 (orange circles), 6, and 7 (open circles) were compared with anti-Ig and CpG-C stimulated cells alone (grey circles). *p<0.05, **p<0.01, ****p<0.0001. The absence of * indicates p>0.05.

### Dynamic CD11c, T-bet, and FcRL5 co-expression profiles in response to stimulation and time

Different combinations of markers are used to describe the B cells that expand during inflammation. Here we analysed how the co-expression of CD11c, T-bet, and FcRL5 change depending on stimulatory conditions at two different time points (**Figure 4**). The largest proportion of triple-positive cells was generated by stimulating the B cells with anti-Ig, CpG-C, and IFN-γ for 2 days (approximately 26%). This condition also had the lowest percentage of cells not expressing any of CD11c, T-bet, or FcRL5 at both 2 and 6 days of stimulation. Compared to 2 days of stimulation, there were few cells expressing all three markers at 6 days. This was mainly explained by the rapid reduction of T-bet^+^ cells after day 2. Instead, the day 6 time-point was characterised by similar frequencies of CD11c single-or CD11c and FcRL5 double-positive B cells. Most of the unstimulated cells remained triple-negative at both time points (88.6% and 76.6% respectively).

**Figure 4.**
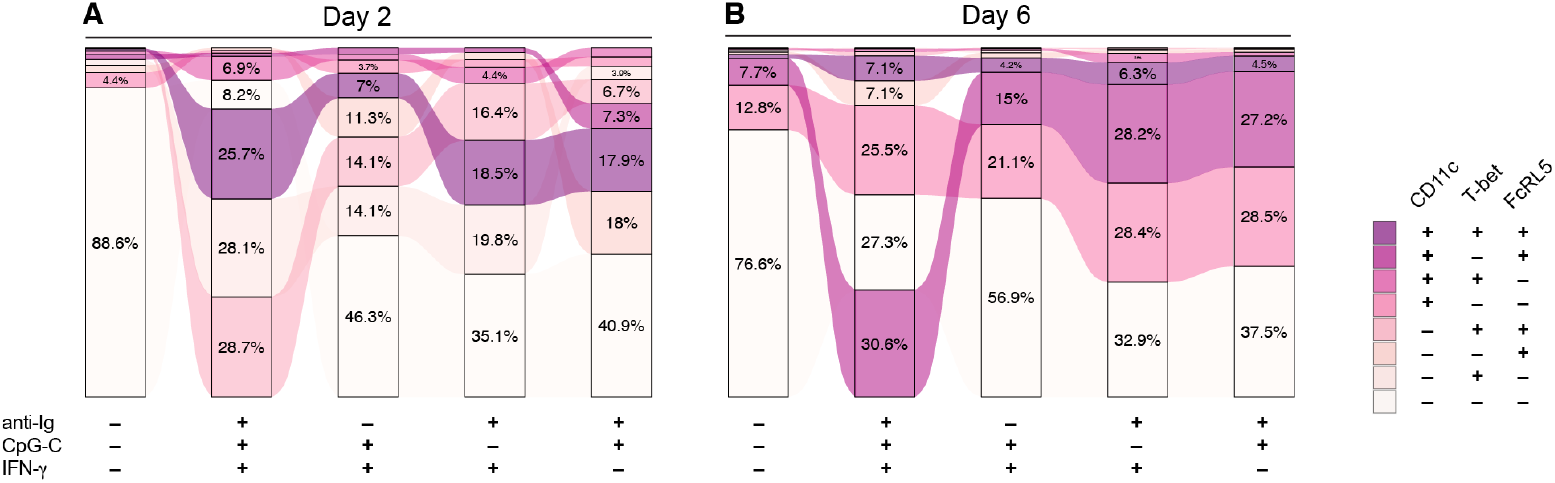
Stimulation-dependent co-expression of CD11c T-bet, and FcRL5. The average frequency of CD11c, T-bet, and FcRL5 co-expression on B cells sorted from buffy coats following stimulation for (**A**) 2 days (n=18) and (**B**) 6 days (n=9) with different combinations of anti-Ig, CpG-C, and IFN-γ. Data is pooled from 7-15 separate experiments.

### Plasma cell differentiation and antibody production are dependent on CpG-C and B cell proliferation

The functional properties of double negative or atypical (CD21^−^CD27^−^IgG^+^ and IgD^−^CD27^−^T-bet^+^) B cells remain unclear. It has been suggested that the cells contribute to the circulating antibody pool (34) and that they could represent an intermediate plasma cell differentiation stage (35). Therefore, we assessed plasma cell differentiation and antibody production in the context of anti-Ig, CpG-C, and IFN-γ stimulation and CD11c, T-bet, and FcRL5 expression. We cultured CTV-labelled and unlabelled B cells sorted from healthy blood donors over 6 days with different combinations of anti-Ig, CpG-C, IFN-γ, and IL-21. We then assessed CTV dilution (indicating proliferation), plasma cell differentiation at 0, 2, 4, and 6 days of culture, and antibody production at 6 days of culture (**Figure 5A**). After subtracting media controls, we compared pooled log2 transformed antibody concentrations (**Figure 5B**). anti-Ig, CpG-C, and IFN-γ stimulated cells (stim. 2) produced significantly more antibodies compared to unstimulated cells (stim. 1). Exchanging IFN-γ with IL-21 (stim. 7) led to further increased antibody levels, although IFN-γ did not have a detrimental effect on the antibody levels (stim. 7 vs stim. 9). Antibody production was however completely abolished in the absence of CpG-C, independent on other stimuli (stim. 4 and 11), indicating that CpG-C was required to plasma cell differentiation.

**Figure 5.**
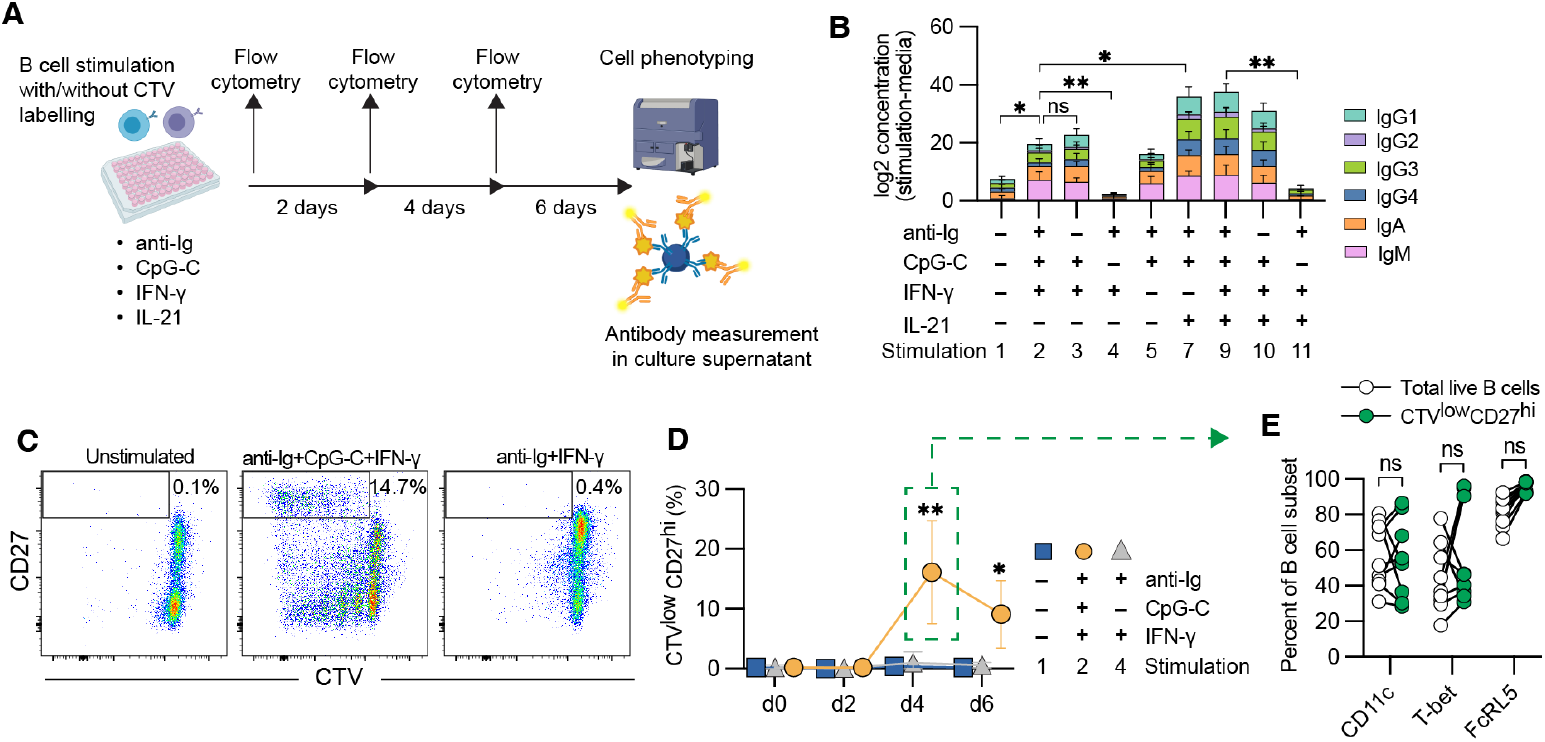
CpG-C is required for plasma cell differentiation and antibody production. (**A**) Schematic illustration of B cell cultures with cell-trace violet labelled and unlabelled B cells sorted from buffy coats, stimulated for 0, 2, 4, and 6 days before (**B**) measuring IgG1, 2, 3, 4, IgA, and IgM antibody levels in culture supernatants associated with different combinations of stimulation with anti-Ig, CpG-C, IFN-γ, and IL-21 as indicated (n=10, pooled from 8 experiments). (**C**) Representative flow cytometry plots from day 6 and (**D**) summary graph indicating the frequency of cell-trace violet low and CD27 high plasma blasts at 0, 2, 4, and 6 days with no stimulation (blue boxes), anti-Ig+CpG-C+IFN-γ (orange circles), or anti-Ig+IFN-γ stimulation (grey triangles) (n=9, pooled from 3 experiments). (**E**) Percent CD11c, T-bet, or FcRL5 expressing B cells within total live B cells (empty circle) or CTV^low^CD27^hi^ cells (green circle) at day 4 after culture with anti-Ig, CpG-C, and IFN-γ. P-values were calculated using repeated-measures one-way ANOVA with Geisser-Greenhouse correction followed by Holm-Sidak’s post hoc test. ns=not significant p>0.05, *p<0.05, ***p<0.001.

We next assessed plasma cell differentiation by flow cytometry. Plasma cells were identified by high levels of CD27, which was only observed in CTV^low^ proliferating cells (**Figure 5C**). Quantifying CTV^low^CD27^hi^ plasma cells after stimulation showed that plasma cells became detectable and reached peak levels at day 4 or stimulation. Plasma cells were, however, only observed after stimulation with anti-Ig, CpG-C, and IFN-γ (stim. 2) and not anti-Ig and IFN-γ alone (stim. 4), or in unstimulated cells. This corroborates the antibody data and shows that CpG-C was critical for B cell proliferation and plasma cell differentiation (**Figure 5D**).

Since it has been indicated that T-bet contributes to plasma cell differentiation (35), we assessed if CD11c, T-bet, or FcRL5 were specifically enriched in plasma cells compared with total live B cells at day 4 of culture, when all markers were expressed and at peak of plasma cell frequencies (**Figure 5E**). Cells expressing either CD11c, T-bet, or FcRL5 were similarly enriched among the two compartments, indicating that none of the proteins was specifically required for plasma cell differentiation (**Figure 5E**).

### B cell stimulation leads to the upregulation of co-stimulatory molecules and cytokine secretion resulting in increased T cell help

In addition to plasma cell differentiation, it has been proposed that CD11c^+^ B cells can provide enhanced antigen-presenting functions to CD4^+^ T cells in mice and humans (19,21). To assess to what extent B cells stimulated with different combinations of anti-Ig, CpG-C, and IFN-γ could drive B cell-dependent CD4^+^ T cell help, we first stimulated B cells from healthy blood donors for 2 days. We then phenotyped the cells, measured cytokines in the supernatants and washed, counted, and co-cultured the B cells at a 1:5 ratio with autologous purified CD4^+^ T cells in the presence of cytostim, crosslinking MHC2 and the TCR (**Figure 6A**). To measure proliferation, the CD4^+^ T cells were labelled with cell trace violet (CTV) (**Figure 6B**). As a negative control, CD4^+^ T cells were cultured with cytostim in the absence of B cells, and as a positive control CD4^+^ T cells were cultured with beads coated with anti-CD3 and anti-CD28. Unstimulated B cells co-cultured with CD4^+^ T cells had a similar frequency of CTV low cells as the negative control, while > 90% of the cells had divided in response to the positive control (**Figure 6B**). B cells pre-stimulated with anti-Ig, CpG-C and IFN-γ (stim. 2) induced significantly more T cell proliferation (CTV^low^ cells) compared with co-cultures with unstimulated B cells. There were also significantly more dividing T cells in the co-cultures with B cells pre-stimulated with anti-Ig and IFN-γ (stim. 4) when compared with unstimulated cells (**Figure 6C**).

**Figure 6.**
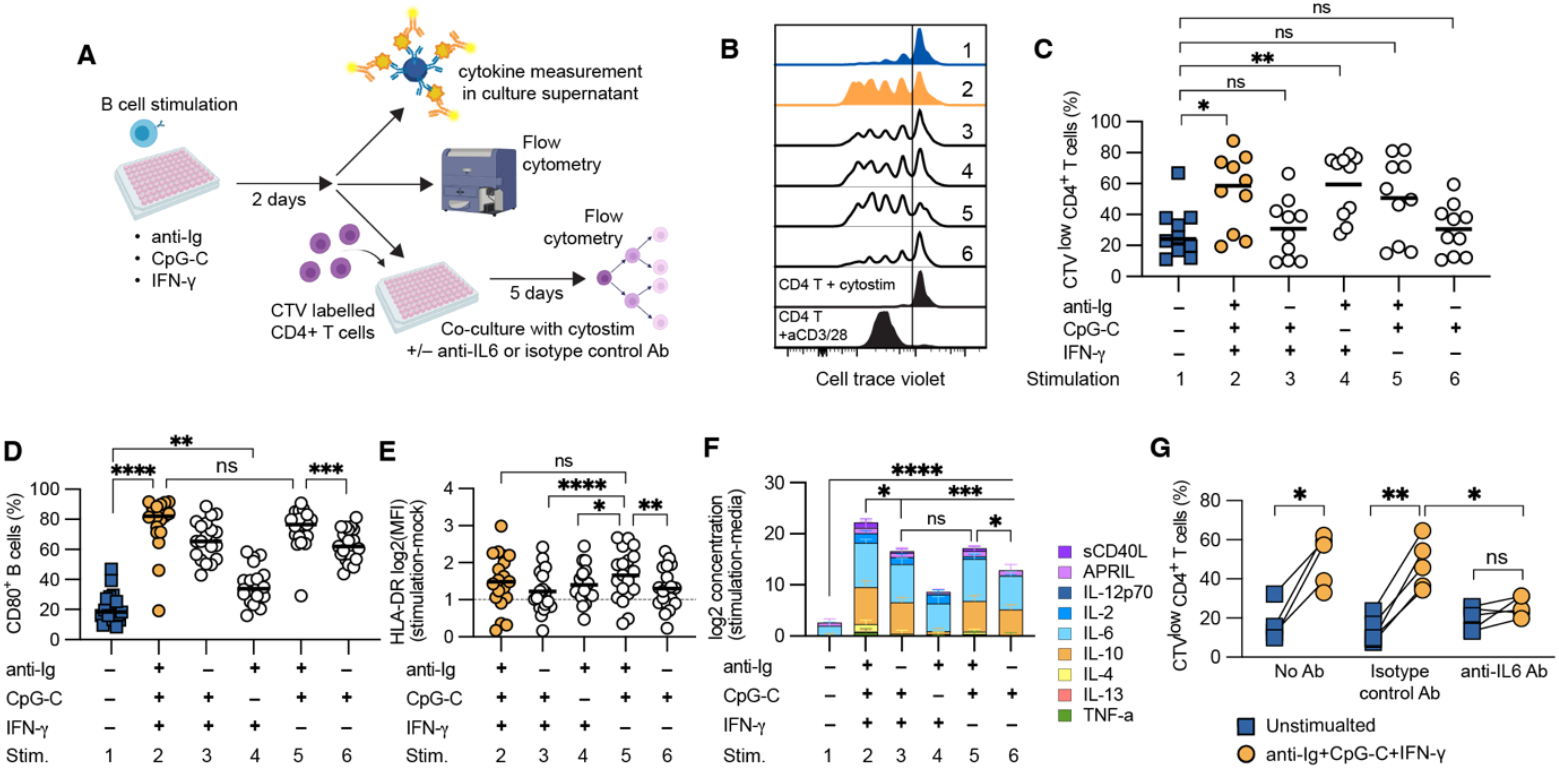
B cell-dependent CD4^+^ T cell proliferation is mediated via IL-6. (**A**) Schematic illustration of B cell-T cell co-culture experiments of cells sorted from buffy coats. (**B**) representative histograms and (**C**) data for cell-trace violet dilution among sorted CD4^+^ T cells co-cultured for 5 days with autologous pre-stimulated B cells in the presence of cytostim (n=10, pooled from 9 experiments). B cells were pre-stimulated for 2 days as indicated, followed by washing, counting, and re-seeding at a 1:5 B:T ratio. (**D**) Frequency of CD80-positive B cells after two days of stimulation (n=19, pooled from 17 experiments). (**E**) B cell surface levels of HLA-DR, calculated as log2[MFI^stim^]–log2[MFI^unstim^], after 2 days of culture (n=19). (**F**) Cytokine levels for sCD40L, APRIL, IL-12p70, IL-2, IL-6, IL-10, IL-4, IL-13, and TNF-? in culture supernatants calculated as log2[pg/mL in stimulated]–log2[pg/mL in culture media] following 2 days of culture (n=10, pooled from 8 experiments). (**G**) The same co-culture setup as in (C) with no stimulation (blue boxes) or anti-Ig+CpG-C+IFN-γ (orange circles) pre-stimulated B cells in the presence of autologous CD4^+^ T cells and No antibody, isotype control antibody, or IL-6-blocking antibody (n=5 from one experiment). P-values were calculated using repeated-measures one-way ANOVA with Geisser-Greenhouse correction followed by Sidak’s post hoc test. For (C), all comparisons were against unstimulated cells. ns=not significant p>0.05, *p<0.05, **p<0.01, ***p<0.001, ****p<0.0001.

In vivo, B cells and CD4^+^ T cells interact via MHC2 and the TCR, with additional survival and polarising signals provided via cytokine secretion and co-stimulatory molecules, such as CD80 and CD86 on the B cells (36,37). To better understand what factors were responsible for the increased B cell-dependent CD4^+^ T cell proliferation, we assessed the levels of CD80 and HLA-DR after stimulation (**Figure 6D**). CD80 was strongly upregulated by anti-Ig, CpG-C, and IFN-γ (stim. 2). The upregulation was independent of IFN-γ as stimulation with anti-Ig and CpG (stim. 5) reached a similar frequency of CD80^+^ cells, while anti-Ig and CpG-C seemed to have an additive effect (**Figure 6D**). All B cells express HLA-DR, so therefore we assessed the surface levels (median fluorescent intensity, MFI) following stimulation after subtracting with the MFI of unstimulated cells (**Figure 6E**). All stimulations that we tested had an average HLA-DR log2(MFI^stim^–MFI^unstim^)>0, indicating upregulation of HLA-DR in response to stimulation. Cells stimulated with anti-Ig and CpG (stim. 5) had a similar HLA-DR MFI level as triple-stimulated cells, while the condition gave significantly more upregulation compared with CpG and IFN-γ (stim. 3), anti-Ig and IFN-γ (stim. 4), or CpG-C alone (stim. 6) (**Figure 6E**).

We next assessed a panel of 13 cytokines in the culture supernatants from day 2 of the B cell culture with the rationale that this could indicate which cytokines were produced by the B cells at the time of co-culture with the T cells. Cytokines that were detected above media background levels (log2[pg/ml^stim^–pg/ml^media^]>0) were pooled together to give a bulk measure of B cell cytokine production (**Figure 6F**). IFN-γ was omitted from the cytokine pool data, since it was spiked in for some stimulations, and is shown as relative concentration (MFI) in **Supplementary Figure 2** together with the other cytokines measured. All stimulations had significantly more cytokines in the culture supernatant compared with unstimulated cells (p<0.0001). anti-Ig, CpG-C, and IFN-γ (stim. 2) stimulated cells had significantly more cytokines than all other conditions (stim. 2 vs stim. 3 p<0.05, stim. 2 vs stim. 4-6 p<0.001). CpG-C + IFN-γ (stim. 3) and anti-Ig + CpG-C (stim. 5) had similar levels of cytokines (p>0.05) and anti-Ig + CpG-C (stim. 5) significantly more than CpG-C alone (stim. 6 p<0.05), indicating that anti-Ig, CpG-C, and IFN-γ all contributed to increasing overall cytokine production (**Figure 6F**). Cells stimulated with anti-Ig and IFN-γ (stim. 4) had the least total cytokine levels of the stimulated cells. This was mainly due to no detectable production of IL-10, which made up 33-39 % of the measured cytokines in the other stimulations, where CpG-C was present. IL-6 was the other main cytokine produced by the B cells, making up 39-64 % of the measured cytokines for the stimulated cells (**Figure 6F**).

To test if IL-6 was potentially important for the improved CD4^+^ T cell help, IL-6 blocking antibodies or isotype control antibodies were added to co-cultures of pre-stimulated B cells and CTV-labelled CD4^+^ T cells. Both the isotype antibody-treated and untreated co-cultures had a significantly higher frequency of dividing CTV^low^ T cells when cultured with anti-Ig, CpG-C, and IFN-γ pre-stimulated B cells compared with unstimulated B cells (**Figure 6G**). Treatment with IL-6 blocking antibodies, however, abolished this effect by significantly reducing the frequency of CTV^low^ T cells to baseline levels, comparable to T cells co-cultured with unstimulated B cells (**Figure 6G**). This indicates that IL-6 was a key factor in the B cell-driven CD4^+^ T cell proliferation.

## Discussion

In this study, we have described how the cell surface proteins CD11c and FcRL5 and the intracellular transcription factor T-bet are dynamically regulated following in vitro stimulation with different activation signals. These proteins are commonly used in different combinations together with CD21, CD27, and IgD to identify subsets of B cells expanding during inflammatory conditions (23).

We have previously shown that many of the dividing B cells during and after acute malaria express CD11c and are enriched for parasite-specific reactivities, indicating a BCR-dependent upregulation of CD11c (27). Similar observations of recently activated B cells being enriched for CD11c^+^ vaccine or infection-specific B cells have also been made with antigen-specific tetramer staining (28,38,39). It is further also consistent with stimulation of the BCR in vitro leading to CD11c upregulation (21). FcRL5 has been shown to be transiently upregulated via BCR-ligation where it upon co-stimulation via the BCR and TLR9 led to a strong proliferation of naïve B cells (40). FcRL5 is an IgG-receptor, enabling B cells to respond to immune complexes (41). In conjunction with CD21 (complement receptor 2), which binds C3-fragments this effect is further enhanced (42), suggesting that early upregulation of FcRL5 could be important to prime B cells for more efficient uptake and subsequent responses to opsonized antigens. CD11c is an integrin and potentially has complementary roles to FcRL5 in enhancing phagocytosis and cell migration (43) that could improve the early B cell response to infection.

During infection, unmethylated DNA from parasites, bacteria, and certain viruses are recognised by monocytes, DCs, and B cells via the expression of endosomal TLR9 and other pattern-recognition receptors (44–47). Binding to the BCR, followed by subsequent antigen uptake into endosomes and signaling via TLR9 constitute early cellular events that the B cells can do independently of other cell-cell interactions. In this study, we found that these events were the primary influence on B cell expression of CD19, FcRL5, CD11c, HLA-DR, CD80, and B cell-derived cytokine production, indicating a role for these effects in the early B cell response to infection. BCR-ligation also leads to the upregulation of CCR7 and EBI2, which will guide the activated B cells to the B cell-T cell interfollicular region in secondary lymphoid organs (48,49), where they can interact with cognate CD4^+^ T cells (49,50).

Prior to their migration to the B cell-T cell interfollicular region, CD4^+^ T cells are primed by activated DCs that have taken up and recognized pathogen associated molecular patterns (PAMPs). In response to intracellular pathogens, DCs produce IL-12, which in turn polarize CD4^+^ T cells to become IFN-γ secreting T_H_1 cells (51). IL-12 together with IFN-γ can then further generate T_FH_1 cells producing both IFN-γ and IL-21 (52). During malaria, circulating T_FH_1 cells, identified by CXCR3-expression are expanded in peripheral blood and are shown to provide suboptimal B cell help compared with conventional CXCR3-negative cT_FH_ cells (53). Especially in children where T_FH_1 cells made up a larger fraction of circulating T_FH_ cells (54). IFN-γ has previously been shown to be important for T-bet expression in human B cells (11) and T_FH_1 cells likely constitute an important driver for the expansion of T-bet^+^ B cells during inflammation (55). Mouse models have in addition to IFN-γ also indicated a role for IL-21 in the generation of CD11c^+^T-bet^+^ B cells (56–58). We similarly show that IFN-γ is critical for optimal T-bet expression but that it is not induced by IL-21 in the absence of IFN-γ. CD11c on the other hand was more strongly upregulated in response to IL-21 than IFN-γ, indicating separate roles of the cytokines and consistent with different regulations of CD11c, FcRL5, and T-bet. In support of this, other studies in mice have identified a population of CD11c^+^ B cells in the absence of T-bet (58,59), and recently, Gao at al. and Gu et al. showed that CD11c is likely to be regulated via the transcription factor Zeb2 in humans and mice, respectively (60,61).

IL-21 is a key cytokine in germinal centre B cell survival and plasma cell differentiation (62). It is mainly produced by CD4^+^ T_FH_ cells and its induction is dependent on interaction with cognate B cells (63). It was shown that CD11c^+^ B cells can become plasma cells in the presence of IL-21 and T cell help (18,21,55,64). Other studies have indicated a contribution of CD21^−^CD27^−^ or CD11c^+^T-bet^+^ B cells to the circulating antibody pool (34,65,66), indicating either active plasma cell differentiation or that the cells are coming from the same lineage as antibody-secreting cells. Further supporting an active plasma cell differentiation, it has been suggested that T-bet^+^ B cells in systemic lupus erythematosus (SLE) could correspond to intermediate-stage plasma cells and that regulation of the IFN-γR by T-bet is important for subsequent full plasma cell differentiation (35). We observed an enhanced effect of IL-21 on antibody production that was unaffected by the presence or absence of IFN-γ in the stimulation and subsequent varying levels of T-bet (lower in the absence of IFN-γ). Additionally, when measuring the expression of CD11c, T-bet, and FcRL5 in plasma cells or other dividing cells responding to the stimulations, there was no enrichment of the markers among plasma cells, suggesting that none of the proteins was actively required for plasma cell differentiation in vitro. However, *in vivo* it is possible that co-expression of these markers is a common occurrence during inflammatory conditions as antigen-specific B cells would upregulate CD11c and FcRL5, after which they would migrate to the interfollicular region where they would encounter T cells producing IFN-γ and IL-21, driving upregulation of T-bet and potentially leading to extrafollicular plasma cell differentiation.

In response to the proinflammatory environment elicited during infections, there is also an expanded anti-inflammatory response characterized by regulatory cells (67,68), modulation of innate immune responses (69–71) and secretion of soluble anti-inflammatory proteins, such as IL-10 (69,72). In mice, an increased level of IL-10 is shown to inhibit T-cell derived IFN-γ, leading to reduced T-bet^+^ B cells (73,74), indicating a potentially dynamic regulation of T-bet^+^ B cells *in vivo*. We also observed a reduced T-bet induction in B cells stimulated with IL-10, but not when co-stimulated with IFN-γ. It has been suggested in mouse and human studies that both B cells and T cells are major sources of IL-10 and are needed for disease control (67,75,76). We observed a strong induction of IL-10 in all stimulations containing CpG-C, consistent with TLR-ligands being major inducers of IL-10 production in B cells (77,78). In addition to IL-10, we also observed a large production of IL-6 from the stimulated B cells. The IL-6 levels were also higher with CpG-C stimulation, but they were not completely CpG-dependent, as they were for IL-10. IL-6 is a pleiotropic cytokine that affects both hematopoietic and non-haematopoietic cells. It is an important survival and differentiation factor for B cells (79) and CD4+ T cells, where it enhances proliferation and influences migration and polarization (80). By blocking IL-6, we completely abrogated the B cell-mediated CD4^+^ T cell proliferation, indicating that IL-6 was a key mediator of how the B cells provided T cell help in our in vitro system.

It has been shown that CD11c^+^ B cells can provide enhanced APC functions in mice (19). We also found that in vitro-stimulated B cells expressing CD11c, T-bet, and FcRL5 could support improved CD4^+^ T cell survival and proliferation via the production of IL-6. However, there was no obvious association with T-bet, as stimulations lacking IFN-γ and therefore were relatively low for T-bet, provided T cell help equally well as stimulation with IFN-γ and high T-bet levels. Instead, the improved T cell proliferation was mainly associated with B cells pre-stimulated via the BCR. This was not clearly associated with specific up-regulation of CD11c, FcRL5 or specific levels of IL-6 either. It is therefore likely that additional factors, driven by BCR-signalling and not identified here, can provide further T cell help in conjunction with IL-6.

B cells expressing CD11c, T-bet, and or FcRL5 remain an enigmatic part of the B cell response. In this study, we have evaluated co-expression after stimulation with different B cell activation signals to learn about how the markers are dynamically regulated over time. We found that the stimuli leading to the upregulation of CD11c and FcRL5 were relatively overlapping, with BCR and TLR9-ligation being the most important, while T-bet was dependent on IFN-γ. We also investigated the stimulated B cells functionally and observed a clear dependency of plasma cell differentiation on proliferation, while T cell help was dependent on the production of IL-6. However, the effector functions were not clearly associated with given patterns of CD11c, T-bet, or FcRL5 expression, potentially indicating that the upregulation of these markers is part of a normal response and not a specific B cell lineage. However, to better understand how these cells behave in humans in vivo, further studies are needed where B cells with different receptor expressions, potentially indicating different stages of differentiation or time after antigen encounter, are characterised for specificity, phenotype, and function.

## Materials and Methods

### Samples – in vitro cultures

For the in vitro parts of the study, cells were sorted from buffy coats from healthy donors. Buffy coats were obtained from the Karolinska University Hospital blood bank, in Stockholm, Sweden. Complete media that was used in all cultures consisted of RPMI-medium supplemented with 10% heat-inactivated FBS, 2mM L-glutamine, 100 U/ml of penicillin, 100 μg/ml of streptomycin, 10 mM HEPES and 0.05 mM 2-Mercaptoethanol (all from ThermoFisher Scientific).

### Total B cell and CD4^+^ T cell isolation

B cells and CD4^+^ T cells were isolated from buffy coats by negative selection with the RosetteSep Human B Cell Enrichment Cocktail or RosetteSep Human CD4^+^ T Cell Enrichment Cocktail respectively, using 50 ml SepMate tubes (all from Stemcell technologies). Briefly, 25 μl of the Cell Enrichment Cocktail was added for each ml of blood and incubated for 20 minutes at RT. Two volumes of PBS + 2% FBS was then added to the sample. 15 ml Ficoll–Paque Plus (GE Healthcare) was added to SepMate tubes and centrifuged at 1000 *g* for 1 minute (RT). Samples were poured into the SepMate tubes and centrifuged at 1200 *g* for 10 min with the break on. The cell fraction was transferred to new tubes and washed twice with PBS + 2% FBS (300 *g*, 10 min, brake at 5, RT). For two donors the B cells were isolated using MACS cell sorting with anti-human CD19 microbeads and MS columns from Miltenyi Biotec, according to the manufacturer’s instructions. After the final wash, the cells were resuspended in PBS + 2% FBS and counted using a Countess II (Invitrogen) following a 1:1 dilution with trypan blue. The cells were centrifuged again and resuspended at 5 × 10^6^ cells/ml in freezing medium (FBS + 10% DMSO) and placed inside a CoolCell at –80°C overnight, after which the cells were transferred to liquid nitrogen for long-term storage. B cell and CD4^+^ T cell purity was assessed by flow cytometry and were on average±SD 90.8±5.0% and 90.5±4.8%, respectively.

### Naïve and memory B cell isolation

Memory B cells and naïve B cells were isolated from RosetteSep pre-sorted total B cells using MACS cell sorting from Miltenyi Biotec. Memory B cells were obtained via positive selection using the Milteny Memory B cell isolation kit, and naïve B cells via negative selection using the Milteny Naïve B cell isolation kit, following the manufacturer’s protocol. Briefly, total B cells were incubated with CD27 Microbeads for 15 minutes before passing through LS columns placed in the MACS separator. The un-labelled fraction was saved for subsequent naïve B cell isolation. The labelled memory B cells were removed from the magnet and eluted. The cells were then counted and resuspended in complete medium before further stimulation or analysis. The un-labelled fraction was incubated with a biotin-antibody cocktail for 5 minutes, followed by another 5 minute incubation with anti-biotin microbeads. The cell suspension was applied to the LS columns placed in the MACs separator and the unlabelled naïve B cells flowing through were counted and resuspended in complete medium before further stimulation or analysis. Memory and Naïve B cell purity were assessed by flow cytometry and were on average±SD 86.7±7.2 % and 83.9±9.2 %, respectively.

### B cell stimulation

Previously sorted total B cells were thawed, moved to a 15 ml Falcon tube and 13 ml of complete media was added dropwise before mixing and resting the cells on ice for 20 min. The sorted B cells were then centrifuged at 300 *g* for 5 min, washed once in complete media (300 *g* 5 min), resuspended in 1 ml complete media and counted. Stimulations of purified B cells were done in a 96-well round-bottom tissue-culture plate (Techno Plastic Products). The stimuli included α-Ig (IgA+IgG+IgM [H+L] AffiniPureF(ab’)2 Fragment goat-anti-human [Jackson ImmunoResearch Laboratories]) at 1.25 μg/ml, CpG-C (Invivogen) at 2.5 μg/ml, IFN-γ (Peprotech) at 25 ng/ml, IL-21 (Peprotech) at 10 ng/ml and IL-10 (Peprotech) at 10 ng/ml. The stimuli were added in different combinations (**Supplementary Table 1**) to wells seeded with 1×10^5^ cells in a total volume of 200 μl complete media. Unstimulated cells were cultured in complete media. Plates were incubated at 37°C and 5% CO_2_.

### Flow cytometry

For flow cytometry, the cells were washed twice in PBS and then incubated with Live/Dead stain (ThermoFisher Scientific), diluted 1:1200 in PBS, for 20 min on ice. Cells were then washed twice in PBS + 2% FBS before incubation for 20 min on ice with an antibody mix targeting cell surface markers. For intracellular markers, the cells were washed once after surface staining, then incubated for 30 min at RT with the FoxP3 Fix/Perm buffer (eBioscience), washed once in FoxP3 Wash/perm buffer, and incubated for 30 min at RT with an antibody mix targeting intracellular markers. The cells were then washed again in Wash/perm buffer followed by one wash in PBS + 2% FBS. Samples were then resuspended in 300 μl PBS + 2% FBS together with 5 μl CountBright™ Absolute Counting Beads (ThermoFisher Scientific) and run on a 5-laser BD Fortessa (BD Biosciences). For the analysis of the naïve and memory B cell, the samples were run on a 5-laser Cytek Aurora (Cytek Biosciences) See **Supplementary Tables 2-8** for the different antibody panels.

### Cell Trace Violet

For tracking cell division, the cells were stained with Cell Trace Violet (CTV) (Thermo Fisher Scientific). The cells were suspended at a density of 2 × 10^6^ cells per ml in PBS. CTV was added to the cell suspension at 1 μM and incubated for 7 min at 37°C in a ventilated incubator. Staining was stopped with FBS (3 times the staining volume) and PBS was added up to 15 ml. The cells were then centrifuged (300 *g* 5 min) and washed once in PBS (300 *g* 5 min) and once in complete media. The cells were then rested for 1 h in complete media at 37°C 5% CO_2_ before further simulation or analysis.

### Antibody measurement

Antibody levels in the supernatants from B cells that had been stimulated for 6 days were measured using the LEGENDplex™ Human Immunoglobulin Isotyping Panel (6-plex) (Biolegend), a bead-base immunoassay, according to the manufacturer’s protocol with some modifications. The assay was performed in a U-bottom 96 well plate, the plate was wrapped in aluminium foil during all incubations. Samples were diluted 1:100 in PBS + 2% FBS. 25 μl diluted sample was mixed with 25 μl assay buffer and 12.5 μl pre-mixed beads. A standard 1:4 serial dilutions were prepared, and 25 μl standard was mixed with 25 μl assay buffer and 12.5 μl pre-mixed beads. The plates were incubated on a plate shaker at 700 rpm for 2 h, RT. Plates were then centrifuged at 650 *g*, 1 min and washed with 200 μl wash buffer. A detection antibody was added at 1:1 in PBS, 25 μl/well. The plates were then incubated on a plate shaker at 700 rpm for 1 h, RT. Streptatividin-PE was then added at 1:1 in PBS, 25 μl/well (no wash in between), and again incubated on a plate shaker at 700 rpm for 30 min, RT. The plates were then washed twice, as before, and resuspended in 200 μl PBS. The fluorescent signal was measured on a BD 4-laser Fortessa using the 488 nm laser (forward scatter vs side scatter) to separate beads based on size and granularity and the 640 nm laser (780/60 filter) to separate beads based on cytokine specificity. The 561 nm laser (582/16) filter was then used to detect the signal, translating to the amount of cytokine. The data was analyzed using the LEGENDplex™ data analysis software.

### Cytokine measurement

Cytokine levels in the supernatants from B cells that had been stimulated for 48 hr were measured using the LEGENDplex™ Human B cell Panel (13-plex) (Biolegend), using the same procedure as described for the antibody measurements, but with the addition of 12.5 μl Matrix B7 to each sample. Supernatants were not diluted prior to being added to the plate and mixed with pre-mixed beads, assay buffer and Matrix B7.

### Co-cultures – in vitro stimulated B cells and CD4^+^ T cells

In vitro stimulated B cells were co-cultured with autologous CD4^+^ T cells and a superantigen to explore the capacity of in vitro stimulated B cells to induce CD4^+^ T cell proliferation. B cells were stimulated in vitro (as previously described) for 48 h. The cells were then washed twice in fresh complete media and counted. 20’000 B cells from each condition were cultured for five days with 100’000 CTV-labelled CD4^+^ T cells along with CytoStim™ (200X dilution, Miltenyi Biotec). TransAct (300X dilution, Miltenyi Biotec) was used as a positive control for CD4^+^ T cell proliferation. To test the hypothesis that IL-6 signalling influences CD4^+^ T cell proliferation, anti-hIL-6-IgG (Invivogen) or mouse IgG1 kappa Isotype Control (eBioscience) were added at 1 μg/ml to the co-culture system at the start, and another 1 μg/ml of anti-hIL-6-IgG (Invivogen) or mouse IgG1 kappa Isotype Control were added to the wells after 2 days of co-culture.

## Data analysis

Statistical analyses and graphs were done using GraphPad Prism (version 9). Data is generally presented as individual data points or mean with error bars indicating the 95% confidence interval. Groups were compared using repeated-measures one-or two-way ANOVA with Geisser-Greenhouse correction of unequal variance or a mixed-effects model if including missing data, followed by Sidak’s test for multiple comparisons, or a Kruskal-Wallis or Friedman test followed by Dunn’s post hoc test for non-parametric data. A p-value < 0.05 was considered significant. Analysis of flow cytometry data was done with FlowJo (version 10.8.1, BD). Alluvial plots were generated using the *ggalluvial* package (81) in R (version 4.1.1, www.r-project.org).

## Data availability statement

Data is available upon reasonable request to the corresponding author.

## Ethics approval statement

This study was performed in accordance with the declaration of Helsinki and following ethical approval (2006/893-31/4 with addendum 2019-03436) by the Swedish Ethical Review Authority.

## Author contributions

LK planned and performed experiments, performed analysis, and wrote the first draft. ADCG planned and performed experiments and wrote parts of the first draft. MJL performed analysis and wrote parts of the first draft. AF supervised students, discussed science, and provided funding. CS planned the study, performed analysis, supervised students, provided funding, and wrote parts of the first draft. All authors contributed to editing the first draft.

## Acknowledgements

We would like to acknowledge the contribution of all the study participants. Many figures were created with Biorender.com. ADCG and MJL were supported by PhD grants from Karolinska Institutet to CS (2020-00878) and to AF (2019-00992), respectively. The research was supported by grants from the Swedish Research Council (2019-01940), the Magnus Bergvall Foundation (2017-02043 and 2018-02656), and the Åke Wiberg Foundation (M18-0076) to CS and by grants from the Stockholm Region (20180409) to AF.

