## Supplemental material for "Regulation of B cell function and expression of CD11c, T-bet, and FcRL5 in response to different activation signals"

**Supplementary table 1. Stimulation matrix**

| Stimulation -> | 1 | 2 | 3 | 4 | 5 | 6 | 7 | 8 | 9 | 10 | 11 |
|---|---|---|---|---|---|---|---|---|---|---|---|
| anti-Ig[1] | - | + | - | + | + | - | + | + | + | - | + |
| CpG-C[2] | - | + | + | - | + | + | + | + | + | + | - |
| IFN-$\gamma$ | - | + | + | + | - | - | - | - | + | + | + |
| IL-21 | - | - | - | - | - | - | + | - | + | + | + |
| IL-10 | - | - | - | - | - | - | - | + | - | - | - |

[1] anti-Ig binds the B cell receptor (BCR)

[2] CpG-C binds toll-like receptor 9 (TLR9)

**Supplementary table 2. Cell purity antibody panel**

| Antibody | Clone | Company |
|---|---|---|
| CD19 PECy7 | HIB19 | BD |
| CD20 APC-H7 | 2H7 | BD |
| CD3 APC-R700 | UCHT1 | BD |
| CD4 BV750 | SK3 | BD |
| CD8 PerCP-Cy5.5 | SK1 | BD |

**Supplementary table 3. B cell antibody panel for the BD LSR Fortessa X-20**

| Antibody | Clone | Company |
|---|---|---|
| CD19 PE-CF594 | HIB19 | BD |
| CD80 BV421 (used on day 2) | L307.4 | BD |
| CD38 BV421 (used on day 6) | HIT2 | BD |
| CD11c BB515 | B-ly6 | BD |
| CD27 BV650 | M-T271 | BD |
| T-bet AF647 | O4-46 | BD |
| HLA-DR APC-H7 | G46-6 | BD |
| FcRL5 PE | 509f6 | Biolegend |
| IgD BB700 | IA6-2 | BD |

**Supplementary table 4. B cell antibody panel for BD LSR Fortessa**

| Antibody | Clone | Company |
|---|---|---|
| IgD BB700 | IA6-2 | BD |
| CD19 PE-Cy7 | HIB19 | BD |
| CD80 BV421 (used on day 2) | L307.4 | BD |
| CD38 BV421 (used on day 6) | HIT2 | BD |
| CD20 BV510 | 2H7 | BD |
| CD11c BB515 | B-ly6 | BD |
| CD27 BV650 | M-T271 | BD |
| CD69 PE-CF594 | FN50 | BD |
| T-bet AF647 | O4-46 | BD |
| HLA-DR APC-H7 | G46-6 | BD |
| FcRL5 PE | 509f6 | Biolegend |

**Supplementary table 5. Antibody panel for cell division and plasma cell differentiation**

| Antibody | Clone | Company |
|---|---|---|
| CD19 PE-Cy7 | HIB19 | BD |
| FcRL5 PE | 509f6 | Biolegend |
| CD27 BV650 | M-T271 | BD |
| CD11c BB515 | B-ly6 | BD |
| CD38 APC-Fire810 | HIT2 | Biolegend |
| T-bet AF647 | O4-46 | BD |
| Cell Trace Violet | | Thermo Fisher Scientific |
| Live/Dead blue | | Thermo Fisher Scientific |

**Supplementary table 6. Antibody panel for assessing B cells in co-culture experiments**

| Antibody | Clone | Company |
|---|---|---|
| CD19 PE-Cy7 | HIB19 | BD |
| CD11c BV650 | B-ly6 | BD |
| CD27 B786 | L128 | BD |
| T-bet AF647 | O4-46 | BD |
| HLA-DR APC-H7 | G46-6 | BD |
| FcRL5 PE | 509f6 | Biolegend |
| IgD BB700 | IA6-2 | BD |
| CD69 PE-CF594 | FN50 | BD |

| | | |
|---|---|---|
| CD80 BV421 | L307.4 | BD |

**Supplementary table 7. Antibody panel for assessing T cells in co-culture experiments**

| **Antibody** | **Clone** | **Company** |
|---|---|---|
| CD3 PE-Cy7 | UCHT1 | BD |
| CD27 B786 | L128 | BD |
| CD62L BV480 | DREG-56 | BD |
| CD45RA APC-H7 | HI100 | BD |
| CX3CR1 AF647 | 2A9-1 | BD |
| CD4 PE | SK3 | BD |
| CD8 PerCP-Cy5.5 | SK1 | BD |
| Cell Trace Violet | | Thermo Fisher Scientific |
| Live/Dead green | | Thermo Fisher Scientific |

**Supplementary table 8. Antibody panel for naïve and memory B cell experiments**

| **Antibody** | **Clone** | **Company** |
|---|---|---|
| T-bet AF647 | O4-46 | BD |
| CD19 APC-R700 | HIB19 | BD |
| FcRL5 PE | 509f6 | Biolegend |
| CD24 PE-CF594 | ML5 | BD |
| CD21 PE-Cy7 | HB5 | Thermo Fisher Scientific |
| CD1c BB515 | F10/21A3 | BD |
| IgD BB700 | IA6-2 | BD |
| CD62L BV480 | DREG-56 | BD |
| CD11c BV650 | B-ly6 | BD |
| CD27 BV786 | M-T271 | BD |
| Cell Trace Violet | | Thermo Fisher Scientific |
| Live/Dead green | | Thermo Fisher Scientific |

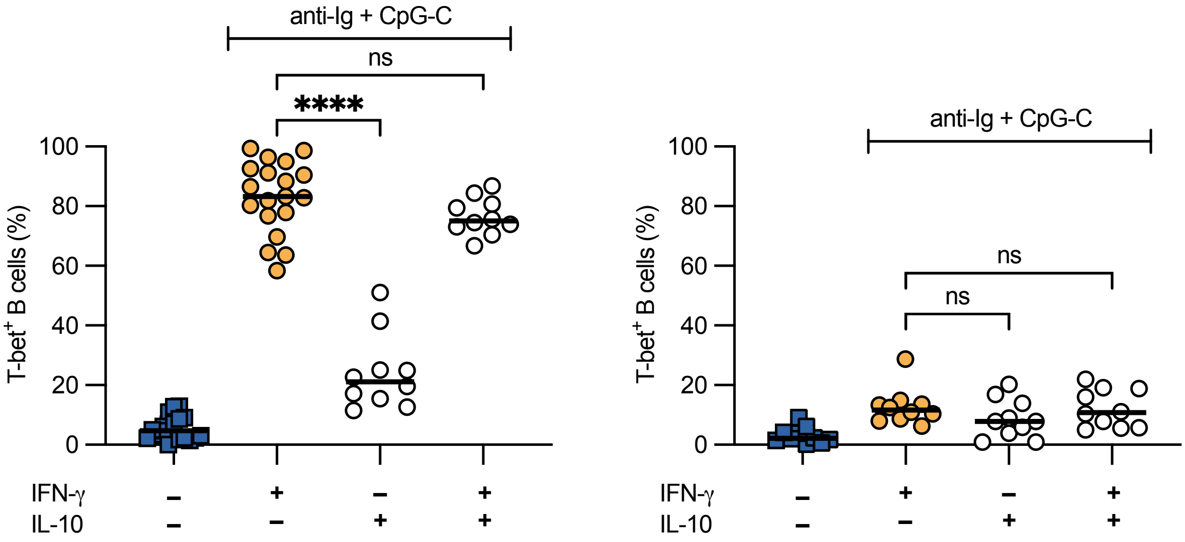


**Supplementary Figure 1. Impact of IL-10 on IFN-γ driven T-bet expression**

Sorted total B cells from buffy coats were stimulated with combinations of anti-Ig, CpG-C, IFN-γ, and IL-10 for (left panel) 2 days (n=11-20) and (right panel) 6 days (n=11) after which expression of T-bet was measured by flow cytometry. Data is pooled from 9-17 separate experiments. P-values were calculated using matched pair one-way ANOVA with Geisser-Greenhouse correction followed by Sidak's posttest or in the case of missing data, by mixed-effects analysis. Unstimulated cells (blue boxes) were included for visual reference. All stimulated groups had received anti-Ig and CpG-C. Cells stimulated with IFN-γ (orange circles) were compared with cells stimulated with IL-10 or IFN-γ+IL-10. ****$p<0.0001$. ns = $p>0.05$.

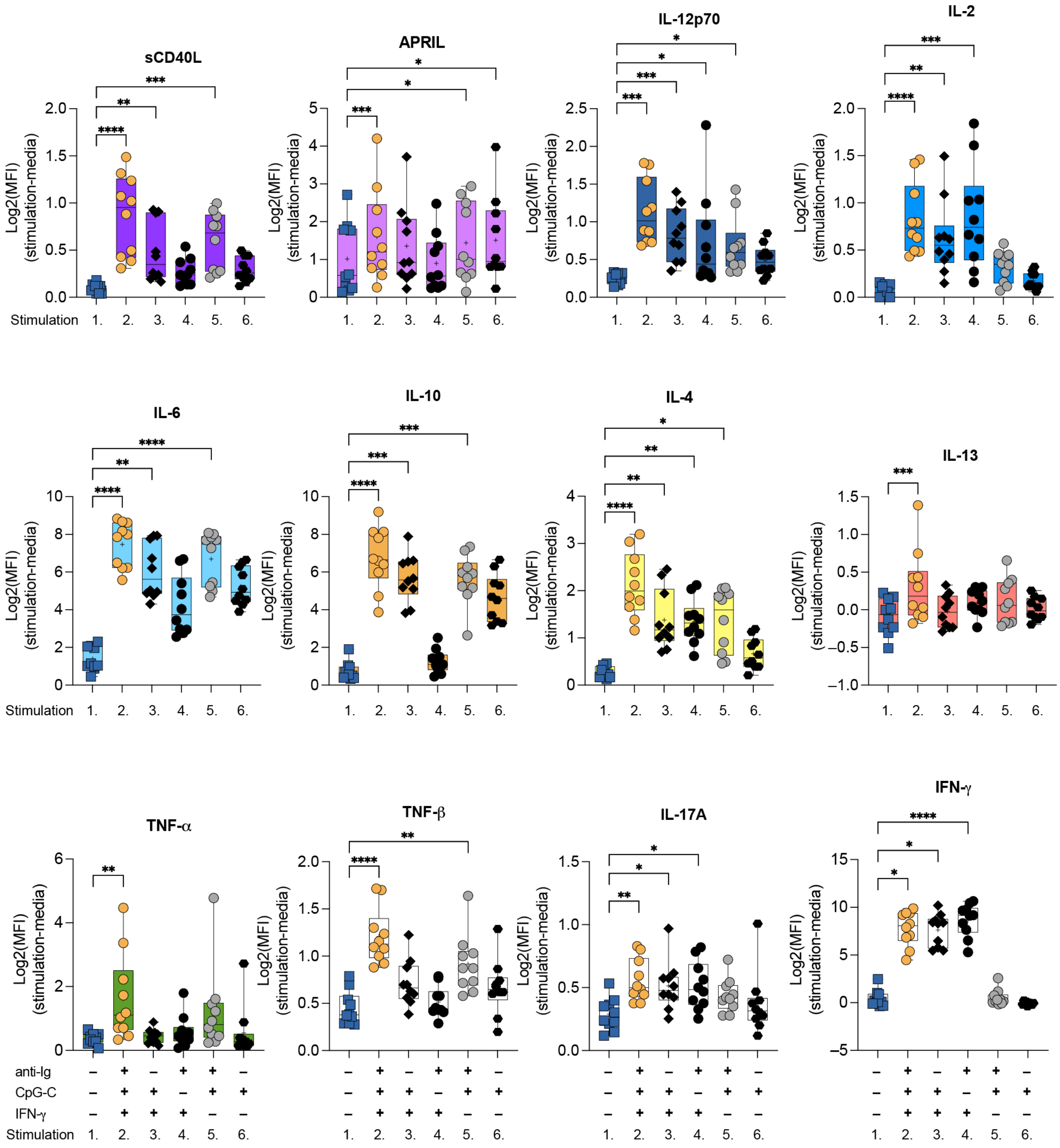


**Supplementary Figure 2. Cytokine levels after 2 days of B cell stimulation.**
A multiplex bead assay was used to measure cytokines secreted by B cells sorted from healthy blood donors (n=10, pooled from 8 experiments) after 2 days of stimulation. The median fluorescent intensity (MFI) for each stimulation, indicating cytokine level, was log-transformed and subtracted from the MFI of the media control. Statistics were calculated using Friedman's test followed by Dunn's posttest to compare all culture conditions with unstimulated cells. *p<0.05, **p<0.01, ***p<0.001, ****p<0.0001. No * indicates p>0.05. The same colour coding is used as for the main figures.
